# Freeze Dried extracts of *Bidens biternata* (Lour.) Merr. And Sheriff. show significant Antidiarrheal activity in ‐vivo Models of Diarrhea

**DOI:** 10.1101/046185

**Authors:** Dennis Gacigi Kinuthia, Anne W. Muriithi, Peter Waweru Mwangi

**Affiliations:** Department of Medical Physiology, School of Medicine, University of Nairobi. P.O. Box 30197-00100, Nairobi, Kenya. Email address; Department of Medical Physiology, School of Medicine, University of Nairobi, P.O. Box 30197-00100, Nairobi, Kenya. Email address

**Keywords:** diarrhea, motility, secretion, jejunum, opioid, adrenoceptor

## Abstract

**Ethnopharmacological relevance of the study:** Diarrhea remains one of the main killers of children aged below five years. Traditional antidiarrheal remedies form a potentially viable source of novel low cost efficacious antidiarrheal remedies in low resource settings. There is therefore a pressing to scientifically evaluate these remedies.

**Aim of the study:** This study aimed to investigate the *in vivo* and *in vitro* antidiarrheal activity of *Bidens biternata* a herb species used in traditional Ayurvedic medicine in the management of diarrhea.

**Materials and Methods:** In the castor oil test twenty (20) adult Sprague-Dawley rats were randomized to the negative control (normal saline), positive control (5 mg/kg loperamide), (200 mg/kg *Bidens biternata* extract) and (400 mg/kg *Bidens biternata* extract) groups (n=5 in each group). Castor oil (4 ml/kg) was then administered to the animals one hour after administration of the respective treatments after which the total mass of fecal output excreted after four (4) hours was determined.

In the charcoal meal test fifteen (15) Sprague Dawley rats were randomized to the negative control (normal saline 5 ml/kg orally), the positive control (atropine sulphate 0.1 mg/kg i.p) and test (400 mg/kg *Bidens biternata* extract) groups (n=5). Charcoal meal was then administered via oral gavage to each rat thirty (30) minutes after the administration of the various treatments. The distance covered by the charcoal meal from the pylorus was then determined after sacrifice of the animals.

In the enteropooling test twenty (20) Sprague-Dawley rats were randomized to the negative control (5% v/v ethanol in normal saline), positive control (5 mg/kg loperamide) and test (400 mg/kg *Bidens biternata* extract) groups and prostaglandin E2 (PGE2) (100μg/kg) administered immediately after the treatments. The animals were then sacrificed half an hour later and the volume of the small intestine contents determined. The effects of different concentrations of *Bidens biternata* extract (0.5. 1.0, 2.0, 3.0 and 5.0 mg/ml) on jejunal contraction were investigated and a dose-response curve constructed using the experimental data after which The ED50 dose determined. The effect of tamsulosin (α1 adrenergic blocker), yohimbine (α2 adrenergic blocker), propranolol (β adrenergic blocker) and naloxone (μ opioid blocker) on the contractile activity of the extract were also investigated.

The experimental data were expressed as mean ± standard error of mean (SEM) and then analyzed using one way ANOVA followed by Tukey’s post hoc test in cases of significance (set at p<0.05).

**Results:** The freeze dried extracts of *Bidens biternata* had significant antidiarrhealeffects in the castor oil induced diarrhea model (p=0.0075) with maximal activity being observed at the 400mg/kg dosage level (1.66± 0.81g vs. 4.54 ± 0.51 g negative control, p=0.01). *Bidens biternata* extract had significant effects on intestinal motility in the charcoal meal test compared to the control group (43.61 ± 4.42% vs. 60.54 ± 3.33%: p= 0.02). *Bidens biternata* extract had a significant effect on PGE2 induced enteropooling (3.06 ± 0.07 ml vs. 4.74 ± 0.10 ml; p<0.001).

The freeze dried extracts of *Bidens biternata* had a significant negative effect on the contractility of the isolated rabbit jejunum (p<0.001). The effects of the extract were significantly attenuated by tamsulosin (53.94 ± 4.20% vs. 80.57 ± 4.09%; p=0.0067) and naloxone (53.94 ± 4.20% vs. 73.89 ± 7.26 %; p=0.0358). Yohimbine (p=0.4598) and propranolol (p=0.5966) however did not have any significant effect on the contractile activity of the extract.

**Conclusions:** The freeze dried extract of *Bidens biternata* possess significant antidiarrhealactivity in both *in vitro* and *in vivo* models which appears to be mediated by modulating both the intestinal motility as well as the secretory activity. The results of this study also validate its traditional use as an antidiarrheal remedy.

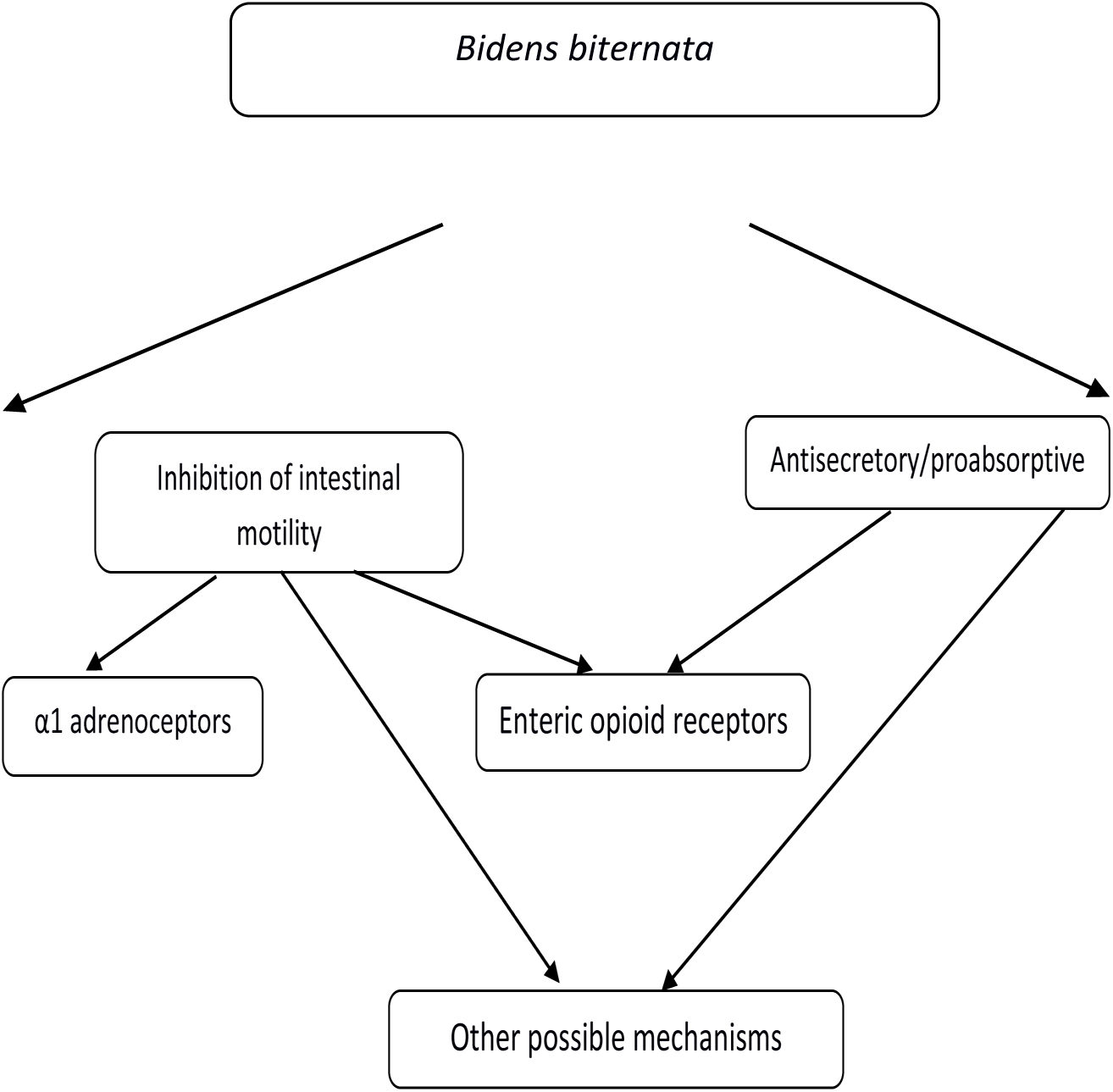

## INTRODUCTION

Diarrhoea continues to be one of the leading causes of death worldwide with areas within the developing world bearing a disproportionate share of the disease burden (Bartolome et al., 2013; Lopez et al., 2006). Indeed it is now documented as the second leading cause of death in children under five accounting for over 1.5 million deaths annually (Kotloff et al., 2013; Walker et al., 2013). Diarrhea is now also recognized also a leading cause of morbidity and mortality among the elderly and HIV infected persons (Balemba et al., 2010; Kosek et al., 2003). There is a therefore a pressing need for the introduction of novel efficacious and less toxic anti-diarrhea treatments.

The World Health Organization states that traditional medicine (TM) is either the mainstay of health care delivery or serves as a complement to it in many parts of the world (Bartolome et al., 2013). There is therefore a pressing need to evaluate traditional medicine therapies for their efficacy as well as possible toxicity.

Plant species from the genus *Bidens* which is a member of the Asteraceae (formerly called Compositae) family are widely used in a variety of traditional medicine systems including Traditional Chinese Medicine(Bo et al., 2012), Aryuvedic medicine (Bartolome et al., 2013; Geissberger and Séquin, 1991)and also in South American traditional medicine (Geissberger and Séquin, 1991; Rabe and van Staden, 1997). Plant species from this genus are often used in the treatment of a variety of gastrointestinal conditions e.g. diarrhea, abdominal pain and even dysentery. Scientific evaluation of a number of plant species has validated their antidiarrheal activity e.g. *Bidens pilosa* L. var. radiata (YADAV et al., 2009), *Bidens odorata*(Zavala-Mendoza et al., 2013) (Astudillo-Valdez et al., 2008). The foregoing discussion explains the rationale for this study which investigated the antidiarrheal activity of this plant species and the potential mechanisms of this activity.

This study aimed at evaluating the efficacy of *Bidens biternata* used in traditional medicine of Paniya and Kattunaayika tribes of Waynadu Districts in Kerala as well in Traditional Chinese medicine the management of diarrhea (Dharmananda, n.d.; Rana and Datt, 1997; Sukumaran et al., 2012).

## MATERIALS AND METHODS

### Plant material and extract preparation

Plant samples containing the aerial parts of *Bidens biternata* were collected from the Naivasha area of Kenya in September 2014. The identity of the collected plant material was verified by resident taxonomists at the Herbarium located at the Department of Botany, School of Biological Sciences, University of Nairobi and a voucher specimen deposited therein (Voucher No. 09202015).

The plant material was air dried for one week after which it was milled to a fine powder using a standard kitchen blender. The resulting powder (500 g) was then macerated with distilled water in a 1:10 (weight: volume) ratio i.e. using five (5) liters of water for half an hour. The resulting suspension was then successively filtered using cotton wool and Whatman’s filter paper respectively. The resulting filtrate was then lyophilized. The freeze dried extract formed was then weighed, placed in amber colored sample bottles and then refrigerated and retrieved as required.

### Animals used

Adult Sprague-Dawley rats aged between six (6) and eight (8) weeks and weighing between two hundred (200) and four hundred (400) g were used in the in vivo studies. Male adult New Zealand white rabbits (N = 6) weighing 2-2.5kg were used in the ex in vivo (isolated jejunum) studies. The animals were housed in the Department of Medical Physiology, University of Nairobi at standard temperature and humidity conditions and 12 hour light/dark cycle. They were fed on food and water ad libitum. They were allowed to acclimatize for 2 weeks before being used for the experiments. In the absence of an Institutional Animal Use and Care Committee (IAUCC), permission for carrying out the study was obtained from the departmental research and postgraduate studies committee. All the experimental procedures used in the study complied with the National Institute of Health (NIH) Guidelines for the care and use of laboratory animals 8th Edition (“Guide for the Care and Use of Laboratory Animals: Eighth Edition | The National Academies Press,” n.d.) and obeyed the 4Rs (reduction, replacement, refinement and rehabilitation) principles in ethical experimental design.

### Castor Oil Induced Diarrhea

The antidiarrheal activity was investigated according to the method described by Zavala-Mendoza (Zavala-Mendoza et al., 2013). Briefly, twenty (20) rats were randomized to the negative control (normal saline), positive control (5 mg/kg loperamide), low dose test (200 mg/kg *Bidens biternata* extract) and high dose test (400 mg/kg *Bidens biternata* extract) groups (n=5 in each group). The treatments were administered by oral gavage. Castor oil (4 ml/kg) was then administered to the animals one hour after administration of the respective treatments also via oral gavage.

The animals were then placed singly in separate cages lined with pre-weighed greaseproof paper. The total mass of fecal output excreted after 4 hours was measured by subtracting the mass of the pre-weighed paper from the total mass of paper and fecal droppings. The results were expressed as a percentage of control according to the model described by Zavala-Mendoza ((Zavala-Mendoza et al., 2013)

### Charcoal meal test

The effect of the extract on normal small intestinal transit was investigated using the charcoal meal method as described by Meite (Meite et al., 2009). Briefly, fifteen (15) Sprague Dawley rats were randomized to the negative control (normal saline 5 ml/kg orally), the positive control (atropine sulphate 0.1 mg/kg i.p) and test (400 mg/kg *Bidens biternata* extract) groups (n=5). The treatments were all administered using oral gavage.

Charcoal meal (10% activated charcoal in 5% gum acacia) (1 ml per rat) was then administered via oral gavage to each rat thirty minutes after the administration of the various treatments. The animals were then sacrificed after half an hour by cervical dislocation, the abdomen cut open and the entire small intestine from the pylorus to the caecum was isolated. The distance covered by the charcoal meal from the pylorus was then measured and expressed as a percentage of the total length of the small intestine.

### Prostaglandin E2 induced enteropooling Test

This test was performed using the method described by Robert (Robert et al., 1976). Briefly twenty (20) Sprague-Dawley rats were randomized to negative control (5% v/v ethanol in normal saline), positive control (5 mg/kg loperamide) and test (400 mg/kg *Bidens biternata* extract). Prostaglandin E2 (PGE2) dissolved in a vehicle consisting of 5% v/v ethanol in normal saline was then administered immediately after the treatments at a dose of 100 μg/kg. The animals were then sacrificed half an hour later by cervical dislocation. The contents of entire small intestine from the pylorus to the caecum were then milked into a beaker and their volume determined.

### Statistical analysis

In all the in vivo studies, the experimental data were expressed as mean ± standard error of mean (SEM). The experimental data was then analyzed statistically using one way ANOVA followed by Tukey’s post hoc test in cases of significance (set at p<0.05) using the Megastat^®^ statistical add-in for Microsoft Excel.

### Isolated Rabbit Jejunum Motility

Five rabbits which had been fasted overnight were used for the study. The rabbits were euthanized by stunning (striking the base of the head in the occipital region) followed by exsanguination. The abdomen was then cut open and a ten (10) cm segment of jejunum isolated and dipped in a beaker containing cold Tyrode's solution (4°C). The contents of the jejunum were flushed out using the Tyrode solution after which the stretch of jejunum was transferred to another beaker containing Tyrode's solution at 37°C. A segment of jejunum about two (2) centimeters long was then suspended in an organ bath containing the Tyrode's solution maintained at 37 ± 1 °C and aerated with carbogen^®^ gas (95% oxygen and 5% carbon dioxide). The upper end of the jejunum segment was hooked to an isometric force transducer (ML500/A, AD instruments) connected to a Powerlab data acquisition system (Powerlab 8/30, AD Instruments)

The jejunum strips were first allowed to equilibrate for an hour before the effects of different concentrations of *Bidens biternata* extract (0.5. 1.0, 2.0, 3.0 and 5.0 mg/ml) on jejunal contraction were investigated. A short record (about 2 min) of normal jejunal activity was recorded before administering each treatment. Starting with the lowest concentration, a one (1) mililiter dose of extract was added to the organ bath and its effect recorded for 3-5 minutes before washout. The organ bath was then rinsed twice with fresh Tyrodes solution and the jejunal strip allowed to regain its baseline activity before administration of the next dose.

The rate and force of jejunal contractions were recorded and analyzed using Chart5^®^ software (AD Instruments). The contraction rate was expressedd as cycles per minute (cpm) while the force of contraction was expressed in grams (g). The effects of the treatments were then expressed in the manner described by Grasa (Grasa et al., n.d.); i.e.activity after treatment/ activity before treatment ^*^100%

And a dose-response curve then constructed. The ED50 dose was then determined from the curve

### Mechanism of Action Experiments

The effect of 1 mg/ml *Bidens biternata* extract on jejunal activity was evaluated in the presence of various standard antagonists (at a concentration of 10^-6^ M); tamsulosin (α1 adrenergic blocker), yohimbine (α2 adrenergic blocker), propranolol (β adrenergic blocker) and naloxone (μ opioid blocker) in an attempt to try and elucidate the possible mechanism(s) of its effects on jejunal activity. Normal jejunal activity was first recorded for two (2) minutes prior to the addition of the antagonist. Jejunal activity after the addition of the antagonist was also recorded further two (2) minutes after which the extract was added and the resulting jejunal activity recorded for a further five (5) minutes before washout. All the experiments were repeated five times.

NB: All the concentrations quoted above were as per the final bath volume (80 ml).

The experimental data were expressed as mean ± standard error of mean (SEM). The experimental data was then analyzed statistically using one way ANOVA followed by Tukey’s post hoc test in cases of significance (set at p<0.05) using the Megastat^®^ statistical add-in for Microsoft Excel.

## RESULTS

### Castor oil induced diarrhea

The freeze dried extracts of *Bidens biternata* showed significant antidiarrheal effects in the castor oil induced diarrhea model (p=0.0075). Post-hoc analysis using the Tukey test showed that the activity was dose dependent with greater activity being observed at the 400mg/kg dosage level (1.66± 0.81g vs. 4.54 ± 0.51 g negative control, p=0.01). However the extract did not possess significant antidiarrheal activity at the 200mg/kg dosage level (p= 0.18). Hence the 400mg/kg dosage of extract was used in the next in vivo experiments. The ED50 was calculated to be 316mg/kg. The experimental data is shown in table 1.

**Table 1.** Inhibitory activity of *Bidens biternata* extract against castor oil induced diarrhea in rats.

| **Treatment Group** | **Dose (mg/kg)** | **Mean mass of fecal output (g) ± S.E.M** | **% Inhibition** |
| --- | --- | --- | --- |
| **Negative Control** | **0** | **4.54 ± 0.51** | **0** |
| **Positive control (Loperamide)** | **5** | **0.72 ± 0.55 \*\*** | **84.16** |
| **Low dose test group** | **200** | **3.13 ± 0.86** | **31.10** |
| **High dose test group** | **400** | **1.66 ± 0.81 \*** | **63.47** |

NS-Normal saline; BBE-freeze dried extract of *Bidens biternata*.
The values are mean ± SEM., n=5. \*p<0.05 and \*\*p<0.01 vs control

### Charcoal meal test

*Bidens biternata* extract had significant effects on intestinal motility in the charcoal meal test compared to the control group (43.61 ± 4.42% vs. 60.54 ± 3.33%: p= 0.02). In addition there were significant differences between the three treatment groups (p=0.02). the experimental data is shown in table 2.

**Table 2.** Effect of *Bidens biternata* extract on small intestinal transit in the charcoal meal test. Key; ^*^p<0.05 and ^**^p<0.01.

| **Treatment** | **Mean**<br>**length of**<br>**small**<br>**intestine**<br>**(cm) ± SEM** | **Mean distance**<br>**travelled by**<br>**charcoal meal**<br>**(cm) ± SEM** | **Mean % movement of charcoal meal**<br>**after 30 min** |
| --- | --- | --- | --- |
| **Negative**<br>**control** | **109.8 ± 6.22** | **66.4 ± 5.30** | **60.54 ± 3.33** |
| **Positive**<br>**control** | **106.1 ± 4.54** | **43.5 ± 7.23** | **40.86 ± 5.54\*\*** |
| **Test** | **113.7 ± 3.34** | **49.1 ± 4.38** | **43.61 ± 4.42\*** |

### PGE2 induced enteropooling

Pretreatment with *Bidens biternata* extract had a significant effect on PGE2 induced enteropooling compared to control (3.06 ± 0.07 ml vs. 4.74 ± 0.10 ml; p<0.001). The experimental data is shown in table 3.

**Table 3.** Effect of *Bidens biternata* extract on PGE2 induced Enteropooling Key ^**^ = p<0.001 and ^***^ = p<0.0001.

| **Treatment**<br>**Group** | **Dose** | **Mean volume of intestinal**<br>**fluid (ml) ± SEM** |
| --- | --- | --- |
| **Negative control** | **100 μg/kg** | **$4.74 \pm 0.10$** |
| **Positive control** | **5 mg/kg** | **$2.68 \pm 0.32^{**}$** |
| **Test** | **400 mg/kg** | **$3.06 \pm 0.07^{***}$** |

### Isolated rabbit jejunum motility

#### Effects of freeze dried extract of *Bidens biternata* (BBE) on spontaneous contractions of the rabbit jejunum

The freeze dried extracts of Bidens biternata had a significant effect on the contractility of the isolated rabbit jejunum (p<0.001). The dose effect on contractility was biphasic with decreases in contractility being observed up to the 3 mg/ml dosage level (42.55 ± 6.26 % of control) after which there was a paradoxical increase in contractility with maximal contractility being observed at 5 mg/ml (61.12 ± 4.30 % of control) dosage level. The experimental data obtained is shown in table 4. The 1mg/ml dose was subsequently used in the mechanism of action experiments.

**Table 4.** The effects of freeze dried extract of *Bidens biternata* on the contraction of the isolated rabbit jejunum. ^*^= p<0.001 and ^**^= p<0.0001.

| BBE<br>(mg/ml) | % of Control (Mean ± SEM) | |
| --- | --- | --- |
|  | Rate of contraction | Force of contraction |
| 0.5 | 94.81 ± 4.95 | 71.01 ± 3.15* |
| 1.0 | 93.52 ± 5.74 | 53.94 ± 4.20** |
| 2.0 | 87.21 ± 7.89 | 45.49 ± 4.81** |
| 3.0 | 93.82 ± 6.60 | 42.55 ± 6.26** |
| 5.0 | 87.25 ± 4.36 | 61.12 ± 4.30** |

#### Effect of antagonists on the inhibitory activity of BBE on the spontaneous contractions of the isolated rabbit jejunum

The negative effects of the extract at the 1.0mg/ml dosage level were significantly attenuated by tamsulosin (53.94 ± 4.20% vs. 80.57 ± 4.09%; p=0.0067%) and naloxone (53.94 ± 4.20% vs. 73.89 ± 7.26 %; p=0.0358). Yohimbine (p=0.4598) and propanolol (p=0.5966) did not have any significant effects on the effects of the extract on the contractility of the isolated jejunum. The experimental data is shown in table 5‥

**Table 5.**
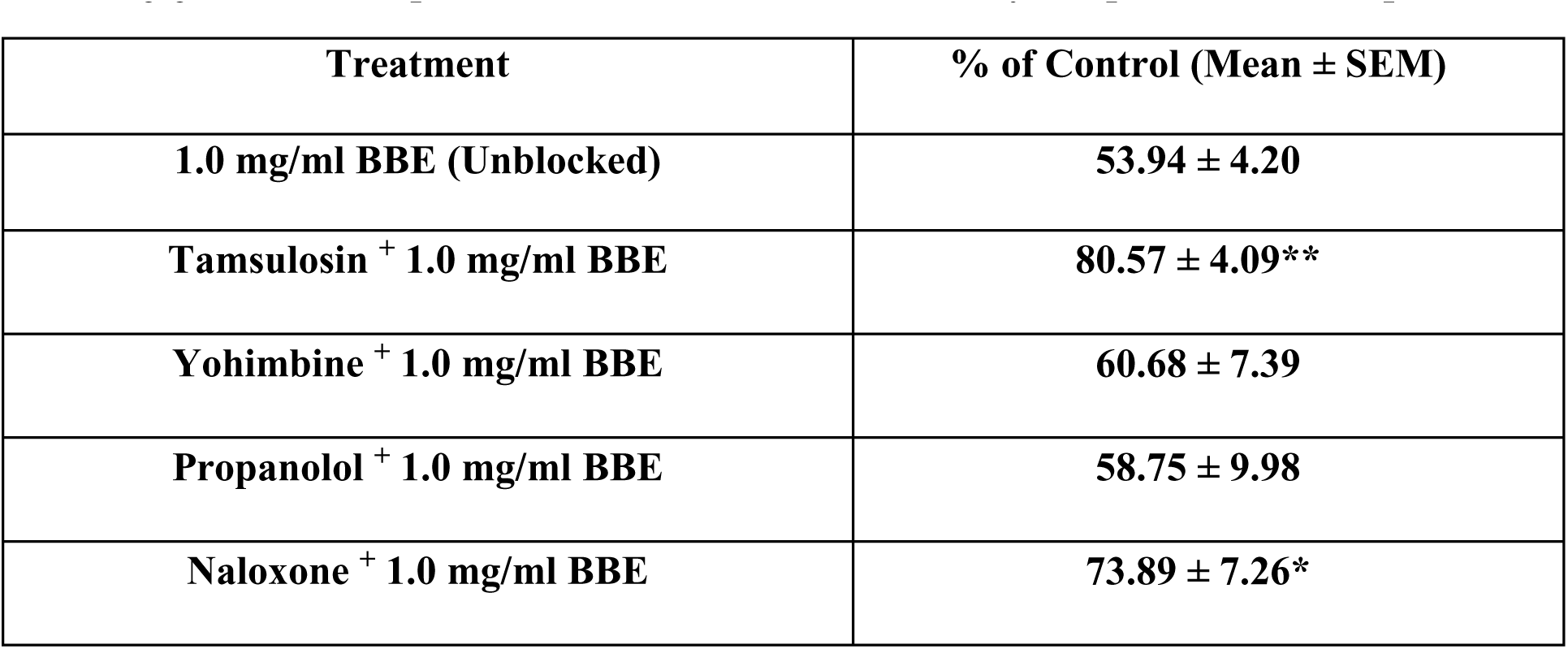
Effects of freeze dried extract of *Bidens biternata* (BBE) on the contraction of isolated jejunum in the presence of various substances. Key; ^*^= p<0.05 and ^**^= p<0.01.

| **Treatment** | **% of Control (Mean ± SEM)** |
| --- | --- |
| **1.0 mg/ml BBE (Unblocked)** | **$53.94 \pm 4.20$** |
| **Tamsulosin + 1.0 mg/ml BBE** | **$80.57 \pm 4.09^{**}$** |
| **Yohimbine + 1.0 mg/ml BBE** | **$60.68 \pm 7.39$** |
| **Propanolol + 1.0 mg/ml BBE** | **$58.75 \pm 9.98$** |
| **Naloxone + 1.0 mg/ml BBE** | **$73.89 \pm 7.26^*$** |

## DISCUSSION

The number of deaths owing to diarrhea remains unacceptably high despite the availability of treatments such oral rehydration salts (ORS), which prevent and treat dehydration, and zinc supplementation, which decreases the severity and duration of diarrhea, and antibiotics for dysentery (Walker et al., 2013). The foregoing discussion underscores the need to discover novel and more efficacious remedies. Herbal remedies which are widely used in the management of diarrhea owing to their affordability, availability and efficacy are a potentially viable source of these new remedies (Sarin and Bafna, n.d.). In addition, herbal remedies also contain multiple chemical constituents which often act synergistically (Gilani and Rahman, 2005; Mehmood et al., 2011).

The freeze dried extract of *Bidens biternata* had significant activity in the castor oil induced diarrhea model which is a model of secretory diarrhea coupled with altered intestinal motility (Zavala-Mendoza et al., 2013). Ricinoleic acid is released by intestinal lipases after oral ingestion of the castor oil (IWAO and TERADA, 1962) induces diarrhea through a number of mechanisms. These include; promotion of Nitric Oxide Synthase activity(Capasso et al., 1986), involvement of endogenous tachykinins (Tiziano *et al.,* 1997), the activation of histamine and serotonin release(Capasso et al., 1986; Gaginella et al., n.d.) among others. Niemegeers (Awouters et al., 1978) suggested that intestinal prostaglandin release was partially involved after they observed that pretreatment with NSAIDs delayed the onset of castor oil induced diarrhea without a significant effect on the fecal output. In addition, ricinoleic acid has been shown to bind to EP3 prostaglandin receptors (Tunaru et al., 2012) and this is then followed by the stimulation of neural reflexes leading to increased motility. The putative mechanisms by which castor oil induces diarrhea are summarized in figure 2. Since the mechanisms by which the extract mediated its antidiarrheal activity were unclear, it was then evaluated in the PGE2 enteropooling assay.

**Figure 1.**
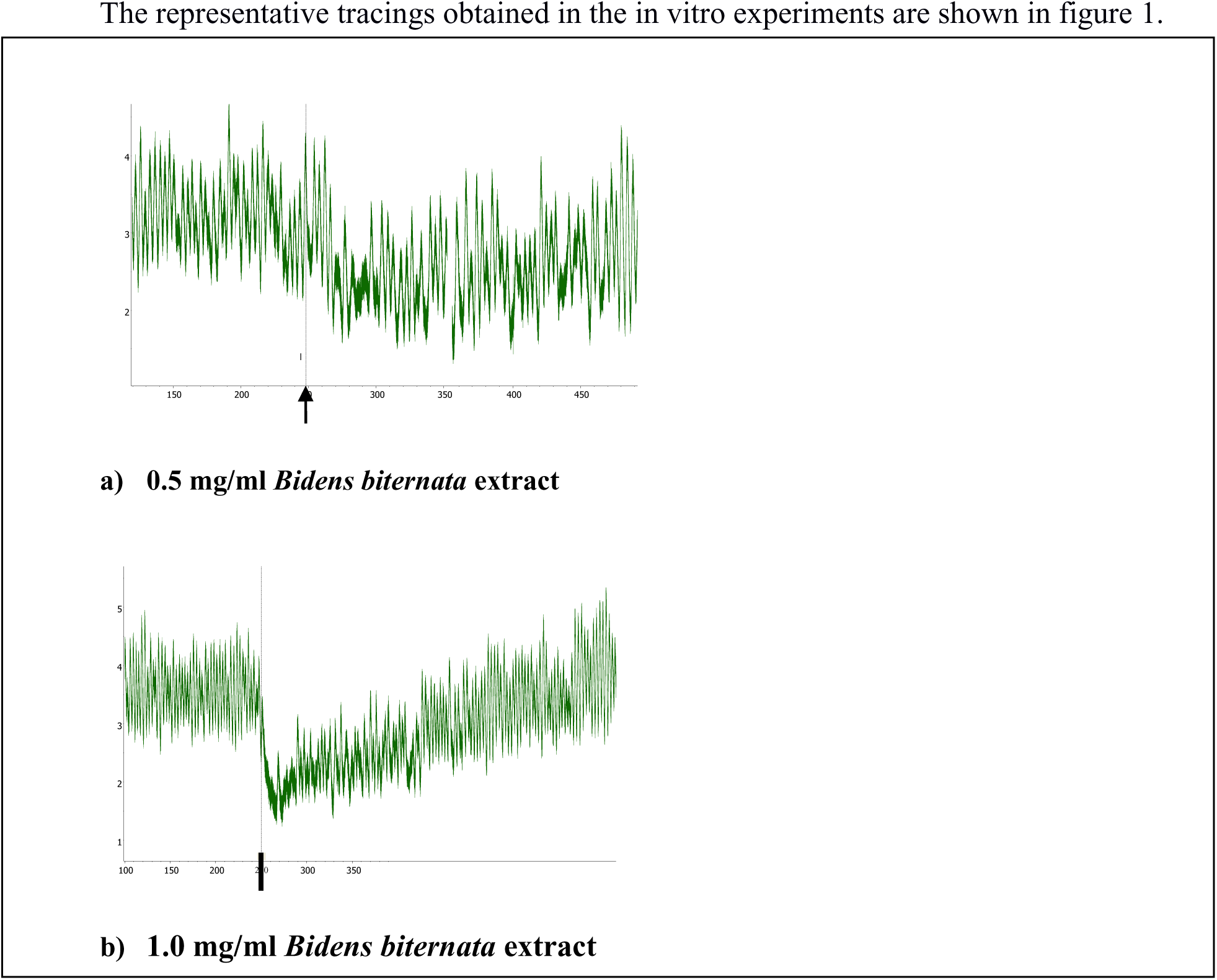
Representative Powerlab tracings obtained in the in-vitro experiments using the isolated jejunum.

**Figure 2:**
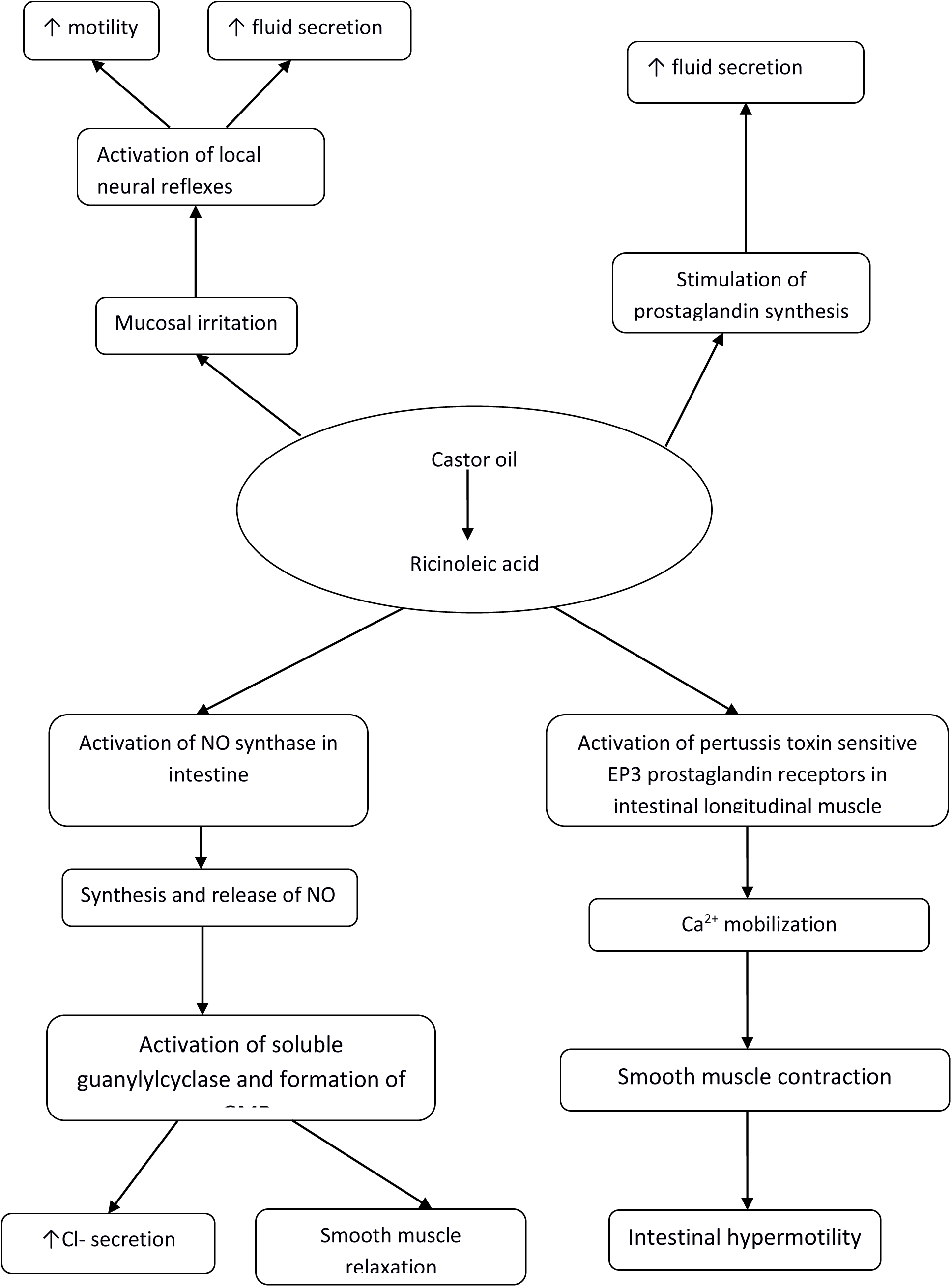
Putative mechanisms of the laxative effects of castor oil

The extract had significant inhibitory effects in the Prostaglandin E2 induced enteropooling model. Prostaglandin E2 is a potent intestinal secretagogue that stimulates cyclic AMP dependent Chloride ion secretion at high doses (Barrett and Keely, 2000; Field, 2003). These effects are mediated via intestinal epithelial EP2 receptors coupled to the Gs family of G proteins (Hosoda et al., 2002; Karaki and Kuwahara, 2004). It can therefore be inferred that the extract contained chemical components that had inhibitory effects on net fluid secretion in the small intestine.

The extract significantly inhibited the movement of charcoal meal. Since charcoal meal is an indigestible substance which distends the intestinal lumen thereby stimulating peristaltic movements (Zavala-Mendoza et al., 2013), this indicates that some of the chemical components of the extract had antimotility effects. The reduction in intestinal motility would potentially allow more time for intestinal fluid and electrolyte reabsorption thereby reducing fluid losses and consequently the fecal output (Sisson, 2011).

It is clear from the foregoing discussion that the extract mediates the observed antidiarrheal effects by modulating both the intestinal secretory and motile activity. The in vitro experiments aimed to elucidate the putative mechanism of action(s) of the observed antimotility activity observed in the vivo models.

The significant antimotility effects of the extract on the isolated rabbit jejunum were attenuated by pretreatment with tamsulosin (α1 adrenergic receptor blocker) and naloxone (nonselective opioid receptor blocker). The antimotility effects were however insensitive to pretreatment with either yohimbine (α2 antagonist) or propanolol (nonselective β adrenergic receptor blocker). it can therefore be deduced that the antimotility activities of the extract are mediated by chemical components of the extract that possess selective agonist activity at α1 adrenoceptors and others that have selective agonist activity at opioid receptors.

It is a well-established fact that the sympathetic nervous system modulates gut muscle motility. α1 adrenoceptors are mainly coupled to the Gq/11 family of G proteins but may also be coupled to the pertussis toxin sensitive Gi protein in certain tissues (Piascik and Perez, 2001). Although α1 receptor agonists trigger smooth muscle contraction in most types of smooth muscle including vascular smooth muscle and sphincters (Garc??a-Sáinz et al., 2000), receptor activation in intestinal smooth muscle causes a decrease in contractility (Rusko and Bauer, 1988).

Activation of the α1receptors causes an increase in the intracellular calcium concentration either by triggering release from intracellular stores via the DAG/IP3 second messenger system or by causing calcium influx into the muscle cells via activation of voltage dependent and voltage independent calcium channels (Piascik and Perez, 2001). The increase in intracellular calcium concentration would ordinarily be expected to lead to contraction. However, it has been shown that there are two calcium distribution compartments in a smooth muscle cell: a contractile compartment centrally located within the cytosol and a non-contractile compartment located between the sarcoplasmic reticulum and the plasmalemma (Karaki et al., 1997; Karaki and Kuwahara, 2004). Agonists that stimulate calcium increment in the contractile compartment trigger smooth muscle contraction while those that trigger calcium increment in the noncontractile compartment trigger other calcium dependent responses such as those related to the cell membrane integrity and permeability (Karaki et al., 1997; Karaki and Kuwahara, 2004). Indeed, Calcium channel blockers and the absence of extracellular calcium have been found to inhibit the α1 mediated effects on intestinal motility, exemplifying the role of calcium in the inhibition of intestinal smooth muscle motility by α1 receptor agonists(Karaki et al., 1997)‥ In addition, the gut inhibitory response to phenylephrine an α1 adrenergic receptor agonist, has been shown to be sensitive to membrane stabilizers such as quinidine and procaine suggesting that its actions may involve alterations in membrane permeability (BOWMAN and HALL,1970). The activity of phenylephrine was on gut motility was shown to be sensitive to apamine (an inhibitor of small conductance calcium-activated channels) (Romanelli et al., 2003) and tetraethylammonium (Rusko and Bauer, 1988). Small conductance calcium-activated channels (SKCa) are voltage-independent potassium channels sensitive to submicromolar concentrations of calcium (Xia et al., 1998). It can therefore be postulated that small increases in intracellular calcium concentration following the activation of the α1 adrenoceptors, would activate these ion channels leading to an efflux of potassium leading to hyperpolarization of the muscle cell and consequently muscle relaxation. This hypothesis would also explain the increase in motility seen at higher doses of extract in the in-vitro experiments.

Naloxone which significantly attenuated the effects of the extract is a nonselective opioid receptor blocker though it has greater affinity for the μ subtype (Holzer, 2009). Opioid receptors are primarily located within the Enteric Nervous System in the gut (Galligan and Akbarali, 2014). Their activation leads to a decrease in neuronal excitability via a variety of mechanisms including;inhibition of adenylyl cyclase, inhibition of Ca^2+^ influx and activation of K^+^ efflux (Al-Hasani and Bruchas, 2011; Galligan and Akbarali, 2014). The reduced excitability in turn causes a reduction in neurotransmitter release in the enteric nervous system e.g. decreased Acetylcholine release from excitatory interneurons, increased NO/purines release from inhibitory motor neurons and decreased Acetylcholine/Vasoactive Intestinal Peptide release from submucosal secretomotor neurons (Galligan and Akbarali, 2014). This then leads to the inhibition of propulsive intestinal motility as well as the inhibition of mucosal chloride ion secretion (Sobczak et al., 2014). It can therefore be postulated that the antidiarrheal actions of the extract may be mediated by binding of some of its chemical constituents to opioid receptors. Although the mechanism of action experiments were carried out using an ex in-vivo model it can be clearly seen that the antisecretory actions of opioid receptor activation can be reasonably extrapolated to explain the antisecretory activity observed in the enteropooling model.

It can then be postulated that the antidiarrheal effects of the extract were mediated by agonism at both the α1 adrenoceptors and the μ opioid receptors. Activity at the α1 adrenoceptors resulted in the inhibition of motility while activity at the opioid receptors was involved in the inhibition of both the motility and secretion.

Preliminary phytochemical analysis of *Bidens biternata* has confirmed the presence of various types of chemical compounds which include phenols, flavonoids, tannins, glycosides and free fatty acids (Sukumaran et al., 2012). In addition, this plant species is used as a leafy vegetable attesting to its nutritional properties which have also been scientifically documented (Sukumaran et al., 2012). This therefore increases its utility in the management of diarrhea that is comorbid with a malnutrition a common situation in the tropics. Future studies will try to identify the specific chemical moieties responsible for the antidiarrheal effects observed in this study.

## CONCLUSIONS

In conclusion, the results obtained in this study indicate that the freeze dried extract of *Bidens biternata* possess significant antidiarrheal activity inin vivo models of diarrhea. The antidiarrheal activity is mediated by the modulating both the intestinal motility as well as the secretory activity. The results of this study also validate its traditional use as an antidiarrheal remedy. Future studies should focus on further evaluation of this plant as a potential source of low cost and accessible antidiarrheal drug moieties

## AUTHOR CONTRIBUTIONS

AWM, PWM, DGK designed the study, DGK did the experimental work, AWM, PWM, DGK analyzed the experimental data and PWM, DGK, PWM wrote the paper. All authors All authors read and approved the final manuscript.

## ACKNOWLEDGEMENTS

The authors wish to acknowledge the great technical assistance provided by Mr David Kayaja Wafula in the execution of his project as well as the technical staff of the department of Medical Physiology, school of Medicine, University of Nairobi. The authors also wish to acknowledge provided by Patrick Chalo Mutiso of the University of Nairobi herbarium for his help in identification of the plants.

## REFERENCES

Al-Hasani, R., Bruchas, M.R., 2011. Molecular mechanisms of opioid receptor-dependent signaling and behavior. Anesthesiology 115, 1363–1381. doi:10.1097/ALN.0b013e318238bba6

Awouters, F., Niemegeers, C.J.E., Lenaerts, F.M., Janssen, P.A.J., 1978. Delay of castor oil diarrhoea in rats: a new way to evaluate inhibitors of prostaglandin biosynthesis. J. Pharm. Pharmacol. 30, 41–45.

Balemba, O.B., Bhattarai, Y., Stenkamp-Strahm, C., Lesakit, M.S.B., Mawe, G.M., 2010. The traditional antidiarrheal remedy, Garcinia buchananii stem bark extract, inhibits propulsive motility and fast synaptic potentials in the guinea pig distal colon. Neurogastroenterol. Motil.: Off. J. Eur. Gastrointest. Motil. Soc. 22, 1332–1339. doi:10.1111/j.1365-2982.2010.01583.x

Barrett, K.E., Keely, S.J., 2000. Chloride secretion by the intestinal epithelium: molecular basis and regulatory aspects. Annu. Rev. Physiol. 62, 535–572. doi:10.1146/annurev.physiol.62.1.535

Bartolome, A.P., Villaseñor, I.M., Yang, W.-C., 2013. Bidens pilosa L. (Asteraceae): Botanical Properties, Traditional Uses, Phytochemistry, and Pharmacology. Evidence-based Complement. Altern. Med. eCAM 2013. doi:10.1155/2013/340215

Bowman, W.C., Hall, M.T., 1970. Inhibition of rabbit intestine mediated by ?‐ and ?‐ adrenoceptors. Br. J. Pharmacol. 38. doi:10.1111/j.1476-5381.1970.tb08528.x

Bo, Y., Yuan, L., Zhang, J., Meng, D., Jing, H., Dai, H., 2012. Total flavonoids of Bidens bipinnata L. a traditional Chinese medicine inhibits the production of inflammatory cytokines of vessel endothelial cells stimulated by sera from Henoch-Schönlein purpura patients. The Journal of pharmacy and pharmacology 64, 882–887. doi:10.1111/j.2042-7158.2012.01480.x

Capasso, F.N.G.V., Mascolo, N., Romano, V., 1986. Laxatives and the production of autacoids by rat colon. Journal of pharmacy and pharmacology 38, 627–629.

Dharmananda, S., n.d. BIDENS.

Field, M., 2003. Intestinal ion transport and the pathophysiology of diarrhea. J. Clin. Investig. 111, 931–943. doi:10.1172/JCI200318326

Gaginella, T.S., Capasso, F., Mascolo, N., Perilli, S., n.d. Castor oil: new lessons from an ancient oil. Phytotherapy Research 12. doi:10.1002/(SICI)1099-1573(1998)12:1+<S128::AID-PTR272>3.0.CO;2-I

Galligan, J.J., Akbarali, H.I., 2014. Molecular Physiology of Enteric Opioid Receptors. Am. J. Gastroenterol. Suppl. 2. doi:10.1038/ajgsup.2014.5

Garc??a-Sáinz, J.A., Vázquez-Prado, J., del Carmen Medina, L., 2000. ? 1-Adrenoceptors: function and phosphorylation. Eur. J. Pharmacol. 389, 1–12.

Geissberger, P., Séquin, U., 1991. Constituents of Bidens pilosa L.: do the components found so far explain the use of this plant in traditional medicine? Acta tropica 48, 251–261. doi:10.1016/0001-706X(91)90013-A

Gilani, A.H., Rahman, A., 2005. Trends in ethnopharmocology. J. Ethnopharmacol. 100, 43–49. doi:10.1016/j.jep.2005.06.001

Grasa, L., Rebollar, E., Arruebo, M.P., Plaza, M.A., Murillo, M.D., n.d. THE ROLE OF NO IN THE CONTRACTILITY OF RABBIT SMALL INTESTINE IN VITRO: EFFECT OF K+ CHANNELS.

Guide for the Care and Use of Laboratory Animals: Eighth Edition | The National Academies Press [WWW Document], n.d. URL http://www.nap.edu/catalog/12910/guide-for-the-care-and-use-of-laboratory-animals-eighth

Holzer, P., 2009. Opioid receptors in the gastrointestinal tract. Regul. Pept. 155, 11–17.

Hosoda, Y., Karaki, S.-I., Shimoda, Y., Kuwahara, A., 2002. Substance P-evoked Cl(-)secretion in guinea pig distal colonic epithelia: interaction with PGE(2). American journal of physiology. Gastrointestinal and liver physiology 283, G347–G356. doi:10.1152/ajpgi.00504.2001

Iwao, I., Terada, Y., 1962. On the mechanism of diarrhea due to castor oil. Jpn. J. Pharmacol. 12, 137–145. doi:10.1254/jjp.12.137

Karaki, H., Ozaki, H., Hori, M., Mitsui-Saito, M., Amano, K.-I., Harada, K.-I., Miyamoto, S., Nakazawa, H., Won, K.-J., Sato, K., 1997. Calcium movements, distribution, and functions in smooth muscle. Pharmacol. Rev. 49, 157–230.

Karaki, S.?. I., Kuwahara, A., 2004. Regulation of intestinal secretion involved in the interaction between neurotransmitters and prostaglandin E2. Neurogastroenterol. & Motil. 16, 96–99.

Kosek, M., Bern, C., Guerrant, R.L., 2003. The global burden of diarrhoeal disease, as estimated from studies published between 1992 and 2000. Bull. World Health Organ. 81, 197–204.

Kotloff, K.L., Nataro, J.P., Blackwelder, W.C., Nasrin, D., Farag, T.H., Panchalingam, S., Wu, Y., Sow, S.O., Sur, D., Breiman, R.F., 2013. Burden and aetiology of diarrhoeal disease in infants and young children in developing countries (the Global Enteric Multicenter Study, GEMS): a prospective, case-control study. Lancet 382, 209–222.

Lopez, A.D., Mathers, C.D., Ezzati, M., Jamison, D.T., Murray, C.J., 2006. Global and regional burden of disease and risk factors, 2001: systematic analysis of population health data. Lancet 367, 1747–1757.

Mehmood, M.H., Siddiqi, H.S., Gilani, A.H., 2011. The antidiarrheal and spasmolytic activities of Phyllanthus emblica are mediated through dual blockade of muscarinic receptors and Ca 2+ channels. J. Ethnopharmacol. 133, 856–865.

Meite, S., N’guessan, J., Bahi, C., Yapi, H., Djaman, A., Guina, F., 2009. Antidiarrheal Activity of the Ethyl Acetate Extract of Morinda morindoides in Rats. Trop. J. Pharm. Res. 8. doi:10.4314/tjpr.v8i3.44533

Piascik, M.T., Perez, D.M., 2001. ?1-Adrenergic receptors: new insights and directions. J. Pharmacol. Exp. Ther. 298, 403–410.

Rabe, T., van Staden, J., 1997. Antibacterial activity of South African plants used for medicinal purposes. J. Ethnopharmacol. 56, 81–87. doi:10.1016/S0378-8741(96)01515-2

Rana, T.S., Datt, B., 1997. ETHNOBOTANICALOBSERVATION AMONGJAUNSARIS OFJAUNSAR-BAWAR, DEHRADUN(U.P), INDIA. Pharm. Biol. 35. doi:10.1080/09251619708951285

Robert, A., Nezamis, J.E., Lancaster, C., Hanchar, A.J., Klepper, M.S., 1976. Enteropooling assay: a test for diarrhea produced by prostaglandins. Prostaglandins 11, 809–828. doi:10.1016/0090-6980(76)90189-1

Romanelli, L., Amico, M.C., Palmery, M., Peluso, I., Savini, G., Tucci, P., Valeri, P., 2003. Role of the cholinergic system and of apamin?sensitive Ca2+?activated K+ channels on rabbit jejunum spontaneous activity and on the inhibitory effects of adrenoceptor agonists. Auton. Autacoid Pharmacol. 23, 105–115.

Rusko, J., Bauer, V., 1988. Calcium and the activation of the alpha 1-adrenoceptors in the guinea-pig taenia caeci. British journal of pharmacology 94, 557–565. doi:10.1111/j.1476-5381.1988.tb11561.x

Sarin, R.V., Bafna, P.A., n.d. Herbal Antidiarrhoeals: A Review.

Sisson, V., 2011. Types of Diarrhea and Management Strategies.

Sobczak, M., Sa?aga, M., Storr, M.A., Fichna, J., 2014. Physiology, signaling, and pharmacology of opioid receptors and their ligands in the gastrointestinal tract: current concepts and future perspectives. J. Gastroenterol. 49, 24–45.

Sukumaran, P., Nair, A.G., Chinmayee, D.M., Mini, I., Sukumaran, S.T., 2012. Phytochemical Investigation of Bidens biternata (Lour.) Merr. and Sheriff.—A Nutrient-Rich Leafy Vegetable from Western Ghats of India. Appl. Biochem. Biotechnol. 167, 1795–1801.

Tunaru, S., Althoff, T.F., Nüsing, R.M., Diener, M., Offermanns, S., 2012. Castor oil induces laxation and uterus contraction via ricinoleic acid activating prostaglandin EP3 receptors. Proc. Natl. Acad. Sci. 109, 9179–9184.

Walker, C.L.F., Rudan, I., Liu, L., Nair, H., Theodoratou, E., Bhutta, Z.A., O’Brien, K.L., Campbell, H., Black, R.E., 2013. Global burden of childhood pneumonia and diarrhoea. Lancet 381, 1405–1416.

Xia, X.M., Fakler, B., Rivard, A., Wayman, G., Johnson-Pais, T., Keen, J.E., Ishii, T., Hirschberg, B., Bond, C.T., Lutsenko, S., Maylie, J., Adelman, J.P., 1998. Mechanism of calcium gating in small-conductance calcium-activated potassium channels. Nature 395, 503–507.

Yadav, A.K., Tangpu, V., Singh, V.K., Govil, J.N., 2009. Therapeutic Efficacy of Bidens pilosa L. var. radiata and Galinsoga parviflora Cav. in Experimentally Induced Diarrhoea in Mice. Phytopharmacology and therapeutic values V 35–45.

Zavala-Mendoza, D., Alarcon-Aguilar, F.J., Pérez-Gutierrez, S., Escobar-Villanueva, M.C., Zavala-Sánchez, M.A., 2013. Composition and Antidiarrheal Activity of Bidens odorata Cav. Evidence-based Complement. Altern. Med. : eCAM 2013. doi:10.1155/2013/170290

